# Bayesian inference captures metabolite-bacteria interactions in a microbial community

**DOI:** 10.1101/2025.09.11.675530

**Authors:** Jack Jansma, Pietro Landi, Cang Hui

## Abstract

Macro-ecosystems, including the human gut, host a vast and diverse set of microbes that indirectly interact with each other through consuming and producing metabolites. Disruptions in this microbial network can affect macro-ecosystem functioning and, in the human gut, contribute to the onset and progression of various disorders, including diabetes, rheumatoid arthritis and Parkinson’s disease. A theoretical foundation for understanding the intricate and dynamic interactions between microbes and metabolites is essential for developing microbiota-targeted interventions to improve macro-ecosystem functioning and health. To this end a precise mathematical framework is crucial to capture and quantify the complex dynamics of the microbial system.

Here, we develop a dynamic network model of coupled ordinary differential equations and present a computational workflow that integrates a generative model with Bayesian inference for model identification. Our approach infers interaction rates, quantifying metabolite consumption and production from simulated time-series data within a Bayesian framework, incorporating prior knowledge and uncertainty quantification. We show that our approach is accurate and reliable in communities of various sizes, sparsity and with different levels of observational noise. This workflow enables *in-silico* predictions of system behaviour under perturbations and offers a robust method to integrate high-dimensional biological data with dynamic network models. By refining our understanding of microbial dynamics, this framework is capable of assessing microbiota-targeted interventions and their potential to improve the health of the macro-ecosystem.

## Introduction

Microbial communities are ubiquitous in nature and play a vital role in all macro-ecosystems such as the soil, ocean, and human gut [1]. Disruption of the delicate balance of microbial communities can be detrimental to the macro-ecosystem. For example, disruption of the soil microbiota alters the regulation of carbon and nutrient cycling, directly impacting macro-ecosystem resilience [1]. Alterations in the composition of the gut microbiota have been implicated in the onset and progression of various disorders including diabetes, rheumatoid arthritis, and Parkinson’s disease [2, 3]. Accordingly, restoration of disturbed microbial communities is a key aspect in the conservation of macro-ecosystems, and in the prevention and treatment of disease. However, a lack of theoretical and computational frameworks of the underlying ecological structure hampers our understanding of microbial communities [4] and subsequently our understanding of the role of these communities in the macro-ecosystem. Developing a theoretical model of microbial communities can support the design of field or wet lab experiments and guide the development of novel interventions that can ultimately improve the health of the host or macro-ecosystems.

Microbial communities are characterised by their composition and abundance but more precisely by their interactions [5]. The majority of these interactions, however, are indirect and mediated via the metabolic environment, specifically via microbial consumption and secretion of metabolites [6]. Therefore, the functioning of a microbial community can be described realistically as the network of these indirect metabolic interactions. Indeed, metabolic concentrations and microbial abundances can be measured experimentally in a community with established methods including metabolomics, sequencing or flow cytometry techniques [7, 8]. However, experimentally measuring the direction and strength of these metabolic interactions is not as straightforward, especially when the interactions are numerous and sporadic, making the quantification of metabolic networks challenging. Most studies use computational methods to infer and quantify microbe-metabolite interactions [9, 10]. For instance, correlations between relative abundances of bacteria across time points (and locations) can serve as edge weights in constructing association network. When combined with bipartite host-microbe networks, these correlation-based methods have revealed distinct nested structures that define the gut microbiota of healthy humans [11]. While correlation-based methods are useful for identifying potential microbe-metabolite interactions, they mainly rely on cross-sectional data and thus overlook the dynamic and context-dependent nature of microbe-metabolite interactions [12].

Adopting a dynamic model not only offers a better representation of the microbiota but paves the way for characterising the stability and resilience of the microbial system and potentially enhances the predictability of microbiota-targeted interventions through scenario simulations. However, longitudinal microbiota studies are costly, limiting the duration and frequency of data collection. For instance, longitudinal sample collection of stool usually takes place in the order of days, whereas the changes within the gut microbiota in response to a perturbation is in the order of hours [13]. Consequently, estimating the frequency and duration of the sampling protocol for longitudinal studies *a priori* becomes essential [13]. Moreover, investigating the dynamics of hard-to-reach microbial communities, such as the small intestinal microbiota, is experimentally challenging [14]. Employing an *in-silico* approach, utilising generative models and statistical inference, can circumvent some of these limitations and better guide the design of field and wet lab experiments, so long as the model itself captures the key dynamic processes of the system.

Dynamical processes in microbial communities are well described using ordinary differential equations (ODEs), and parameter inference is well established using frequentist methods such as least squares [15]. However, these methods usually require large amounts of data and specific assumptions about the residual distribution to provide reliable estimates. When data are sparse, as is common in high-dimensional, nonlinear systems such as in microbiota dynamics, least-squares estimation can become unstable, prone to overfitting, and may underestimate uncertainty, leading to misleading point estimates. In addition, it offers no formal way to incorporate prior knowledge [16]. In contrast, Bayesian inference naturally accounts for uncertainty in the data through likelihood functions and incorporates prior information, yielding posterior distributions from which credible intervals and parameter correlations can be derived. It also enables direct assessment of whether the inferred model reproduces observed data patterns. Moreover, Bayesian methods are well suited for nonlinear models and, by combining priors with the likelihood, can stabilise parameter estimation and address issues of identifiability (i.e., different combinations of parameters can explain the data equally well, making it difficult to distinguish among them) [17].

Here, we develop a bacteria-metabolite model using a set of ODEs, describing the consumption and excretion of metabolites by bacteria in a community. Using simulated data from this generative model, we employ Bayesian inference to quantify bacterial-metabolite interactions in bacterial communities of various sizes and sparsity. Using Bayesian inference to address model identifiability, further allows us to inform the design of longitudinal experiments for optimised sampling strategies. The combination of a generative mathematical model with a Bayesian statistical approach, therefore, contributes to our fundamental understanding of microbial community functioning.

## Material and methods

### Model construction

Our model is defined by a set of coupled ODEs that describe the dynamics of bacteria and metabolites in a community [18]. Let *M*_*i*_ depict the concentration of metabolite *i* and *B*_*j*_ the abundance of bacterium *j* in a microbial community, where all bacteria solely interact indirectly via consuming and excreting metabolites in the shared metabolic environment (denoted by vector ***M***= {*M*_1_, *M*_2_,… }). After considering the death rate *d*_*j*_ of bacterium *j* in a finite environment with carrying capacity *k*, we can consider the per-capita growth rate, *g*_*j*_(***M***), that depends on the uptake and assimilation of multiple metabolites.

Consider for example the simplest form, where assimilated metabolites are substitutable and additively contribute to the per-capita growth rate, 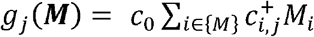, where *c*_0_ depicts the metabolite conversion coefficient and describes how much of the total consumed metabolites is converted into population growth, and *c*_*i*,*j*_ depict the consumption (*c*_*i*,*j* >_0) or excretion (*c*_*i*,*j*_0) rate of metabolite *i* by bacterium *j*, with 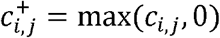 representing the fact that only consumed metabolites support cellular processes and contribute to bacterial population growth. Metabolites secreted into the environment are considered waste products that do not affect bacterial population growth. This yields the following population dynamics of the bacteria:

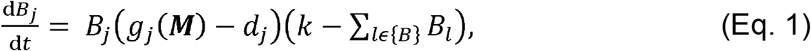

where {*B*}is the set of bacteria species. We assume that the consumption of metabolites is sufficiently fast to deem the term for metabolic outflow or degradation unnecessary. The dynamics of the metabolites are therefore defined as:

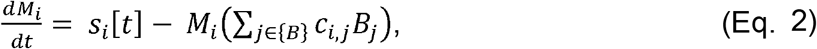

where *s*_*i*_[*t*] depicts the supply rate of metabolite *i* at time *t*, with *s*_*i*_[*t*] =0 representing a closed system where the initial metabolite concentration *M*_*i*_[0] can only increase from bacterial secretion (*c*_*i*,*j*_0) or decline from bacterial consumption (*c*_*i*,*j*_ > 0). Note, the system becomes open if *s*_*i*_[*t*] ≠ 0, which can reflect the manipulation of the metabolic environment.

For the purpose of this study, we fixed the death rate, *d*_*j*_, at 0.1, the metabolite conversion coefficient, *c*_0_, of each bacterium at 0.1 [19, 20] and the carrying capacity at 14, which is approximated and scaled from the total number of bacterial cells (14 billion per mL) obtained in a wet lab experiment [21]. We also fixed *s*_*i*_[*t*] =0 in Eq. (2) to represent the closed system in this wet lab experiment. Consequently, the focus below is to estimate the interaction coefficients, *c*_*i*,*j*_, using count and concentration readings of *B*_*j*_ and *M*_*i*_, respectively, from snapshots of the microbiota at different time points.

To generate synthetic data of modelled microbiota dynamics, we randomly assigned interaction coefficients, *c*_*i*,*j*_, and initial conditions (*M*_*i*_[0], *B*_*j*_[0]) and solved the model using the Runge–Kutta 4(5) solver (`ode45`) in the R package `deSolve`. Let the observations of the quantity of bacterial or metabolic species, *l*, at timepoint, *t*, in replicate, *r*, be denoted as, *y*_obs_ = {*y*_*r*,*t*,*l*_}. Note, the observed value *y*_*r*,*t*,*l*_ and the predicted value *µ*_*r*,*t*,*l*_ from the model (Eq.1&2, representing the assumed microbiota dynamics) are both non-negative, and they can be different due to the observation error *ε*_*r*,*t*,*l*_ defined on the logarithmic scale,

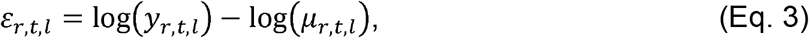

where in practice a small positive constant δ close to zero is added in the logarithm to avoid zero quantity. Observation errors can arise due to sample handling and measurement error in real world experiments. The observation error is assumed to follow a normal distribution with zero mean and standard deviation σ_err_. Consequently, the simulated observations can be generated from a log-normal distribution with the predicted value as the mean:

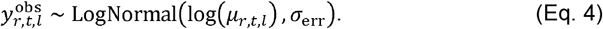

First, we generated the dynamics of a small community, without observation error 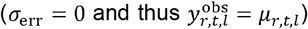, consisting of three bacteria, three produced and seven consumed metabolites, with their initial values randomly assigned from Uniform(0.05, 0.2), Uniform(*e*^-3^, *e*^-2^) and Uniform(0.5, 1), respectively, while the interaction coefficients of the produced metabolites 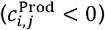 and the consumed metabolites 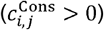 were randomly obtained from Uniform(-1, 0) and Uniform(0, 1), respectively. For each set of randomly assigned interaction coefficients, we generated six independent replicates using different initial conditions. The generated dynamics (*µ*_*r*,*t*,*l*_) were observed at four evenly spaced timepoints between *t*= 0 and *t*= 4. Second, to assess the effect of observation error, we computed the observed value 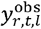 in Eq. 4 with the error standard deviation, σ_err_, independently drawn from Uniform(0.1, 0.5) for metabolite concentrations and from Uniform(0.01, 0.05) for bacterial densities, for each actual snapshot value, *µ*_*r*,*t*,*l*_, of the generated dynamics.

### Bayesian inference

In the Bayesian inference framework, the posterior distribution, P (*c*_*i*,*j*_, σ_res |_ *y*_obs_), of interaction coefficients, *c*_*i*,*j*_, and residual standard deviation, σ_res_, is described by:

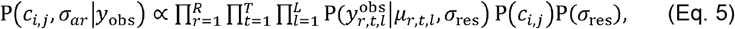

where 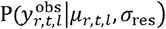 is the likelihood of observing 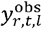 given the model prediction *µ*_*r*,*t*,*l*_ and residual standard deviation, σ_res_; P (*c*_*i*,*j*_) and P(σ_res_) depict the priors. To assess whether the Bayesian inference framework could correctly estimate the assigned interaction coefficients from simulated observations, we specified weakly informative priors to regularise estimation while retaining flexibility. Interaction coefficients were assigned according to their metabolite category: 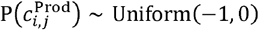∼ Uniform(-1, 0) for produced metabolites and 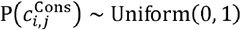 ∼ Uniform(0, 1) for consumed metabolites. The prior for the residual standard deviation was specified as 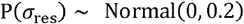 ∼ Normal(0, 0.2).

Bayesian inference was performed using Hamiltonian Monte Carlo with the No-U-Turn Sampler (NUTS), as implemented in the R package `Rstan` [22]. For simulated data without observation errors, inference was conducted with four chains of 2000 iterations, discarding the first 1000 iterations as warm-up. To reduce computational costs for all other simulated data with randomly assigned observation error, as well as for larger and sparser communities, two chains of 1000 iterations were run, with 500 warm-up iterations discarded. To improve sampling performance in the presence of complex posterior geometries, we set the target acceptance probability (`adapt_delta`) to 0.99, thereby reducing the risk of divergent transitions.

The quality of inference was evaluated through standard convergence diagnostics and posterior predictive checks. Convergence was assessed with the potential scale reduction statistic (`Rhat`), while sampling efficiency was quantified as the effective sample size (`n_eff`). Posterior predictive checks were performed by comparing the input observations with simulated observations obtained by solving the ODEs using the `integrate_ode_rk45`, as implemented in the R package `Rstan`, with interaction coefficients drawn from the posterior. Model accuracy was further assessed by calculating the mean absolute error (MAE) between the assigned interaction coefficients and their posterior means, as well as by calculating the coverage probability, which gives the proportion of assigned interaction coefficients that fall within the 95% credible intervals of the posterior distribution.

## Results

### Accurate and reliable estimation of interaction coefficients

For communities without observation errors, model diagnostics of the Bayesian inference framework indicated proper chain mixing **(Figure 1A)**, no divergent transitions, with all `Rhat` values below 1.002 and all `n_eff` above 5500, collectively indicating sampler convergence and reliable parameter estimation across all model parameters **(Figures S1A & B)**. Inspecting the posterior distributions showed that the credible interval is small and that the means are spread out across the full range of the assigned priors **(Figure 1B)**. Posterior predictive checks showed that solving the set of ODEs with interaction coefficients drawn from the posterior distributions produced predictions consistent with the observations **(Figure 1C)**, with posterior means overlapping well with the assigned interaction coefficients **(Figure 1D)**. Overall, Bayesian inference showed robust inferential performance and high parameter precision, underscoring methodological accuracy.

**Figure 1:**
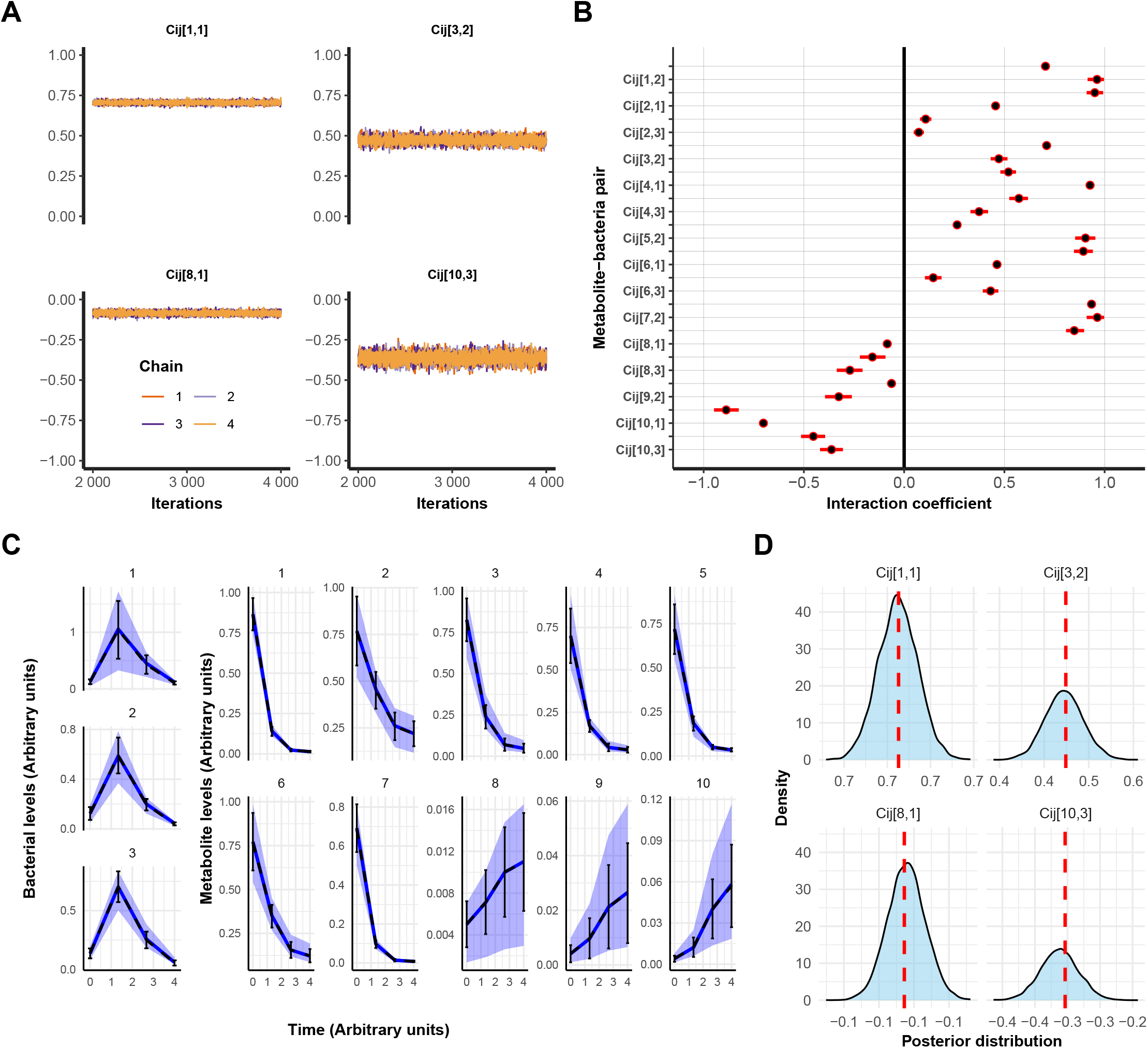
Simulation-based calibration accurately estimates interaction coefficients in a small bacterial community. A) Trace plots of the posterior samples for four representative metabolite-bacteria interaction coefficients across four independent Hamiliton Monte Carlo-NUTS chains. Each panel shows sampled parameter values over iterations. B) Posterior distributions of all interaction coefficients, *c*_*i*,*j*_. Posterior means are depicted as black dots, with 95% credible intervals indicated by red bars. C) Posterior predictive check. The mean and standard deviation of four generated observations, obtained by numerically solving the set of ODEs using the assigned initial conditions and interaction coefficients, are shown as black dotted lines and error bars. The mean and 95% credible intervals of generated observations, using draws from the estimated posterior distributions, are shown as a blue line and a purple belt. D) Four representative comparisons of assigned interaction coefficients (red dotted lines) with the corresponding posterior distributions.

We evaluated the reliability of our inference approach by comparing the assigned interaction coefficients with the corresponding posterior mean across 250 independent simulations. The median of the MAE was 0.027, with 75% of simulations below 0.04, indicating consistent recovery of interaction coefficients **(Figure 2A)**. Next, we evaluated the accuracy by comparing every assigned interaction coefficient, *c*_*i*,*j*_, with the corresponding posterior mean for all simulations, resulting in a total of 7500 comparisons **(Figure 2B)**. The median of these variable-specific comparisons was 0.018, with 75% of simulations below 0.04, with the maximum error obtained being 0.565 **(Figure 2C)**. Lastly, we calculated the coverage probability, representing the proportion of assigned interaction coefficients that fell within the 95% credible interval of their corresponding posterior distributions, indicating high reliability of our inference approach **(Figure 2D)**.

**Figure 2:**
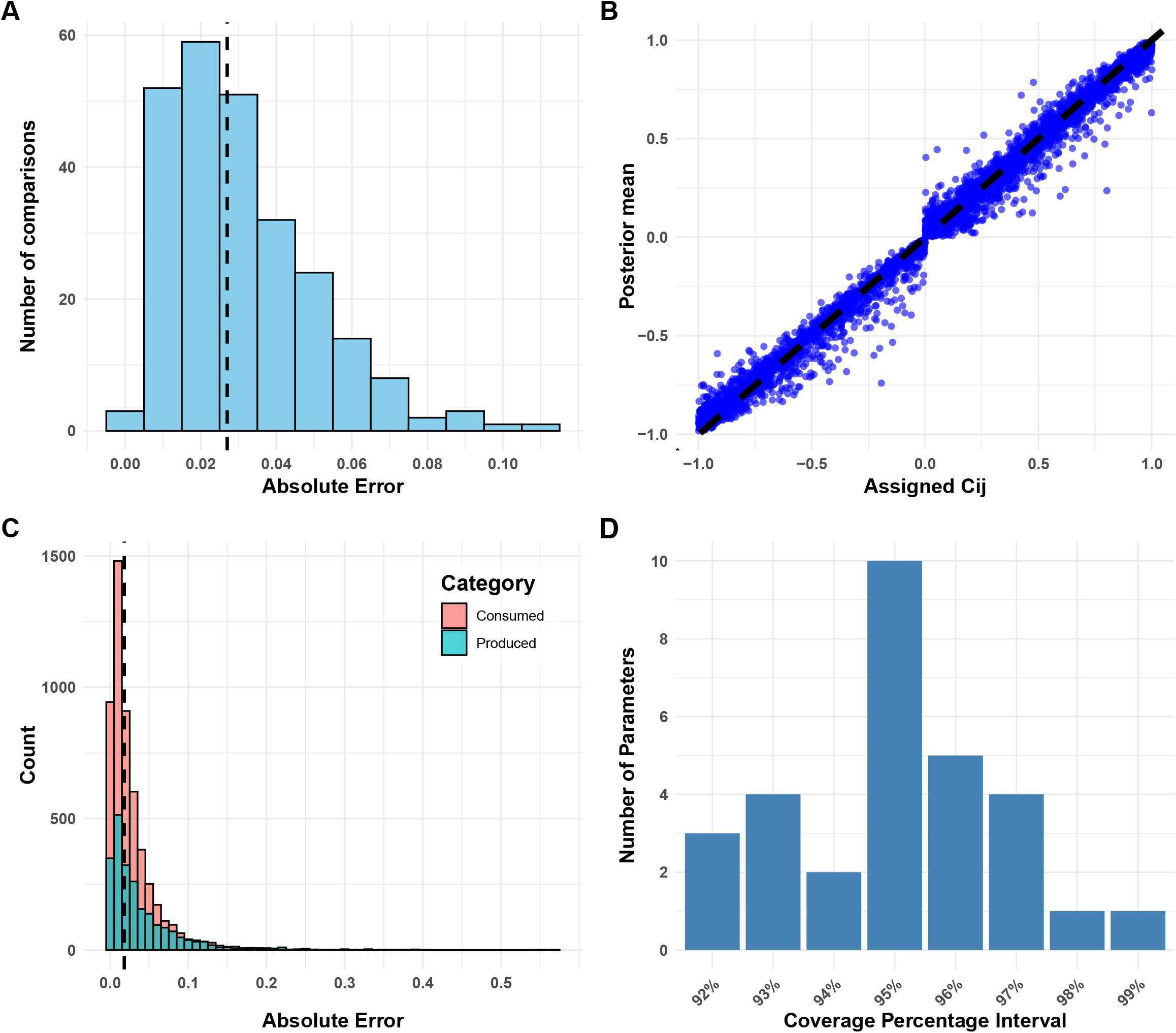
Independent simulations show reliable estimation of interaction coefficients in a small bacterial community. A) Mean absolute differences between assigned interaction coefficients and corresponding posterior means across 250 simulations. The median is shown as the black dotted line. B) 7500 comparisons of assigned versus estimated parameters across 250 simulations. The x-axis represents the assigned interaction coefficients (*c*_*i*,*j*_), and the y-axis depicts the corresponding posterior mean. The black dotted line indicates the 1:1 relationship expected if posterior means matched the assigned values exactly. C) Absolute differences between assigned interaction coefficients and corresponding posterior means, plotted across 7500 parameters across 250 simulations. Interaction coefficients for produced and consumed metabolites are shown in green and red, respectively. The median is shown as the black dotted line. D) Distribution of the percentage of assigned interaction coefficients that fall within the 95% credible interval of the posterior distribution across 250 simulations.

### Reduced uncertainty with increased number of observations

The addition of observation errors did not negatively affect the diagnostics of the Bayesian inference framework **(Figures S2A & B)** although more uncertainty was found as indicated by broader posterior distributions **(Figure 3A)**, which was also reflected in the posterior predictive check **(Figure S2C)**. A straightforward way to increase the accuracy of parameter estimation is to increase the number of observations. We tested this hypothesis by increasing the number of observations of the small bacterial communities under random observation errors from 4 observations to 6, 8, 10 and 25, evenly spaced across the same temporal range. Model diagnostics was not affected by increased observations **(Figures S3A & B; S4A & B)** but the credible interval of the posterior distributions was reduced **(Figure 3B)**, with the smallest credible interval obtained using the largest number of observations **(Figure 3C)**. Furthermore, a posterior predictive check indicated accurate recovery of the observed data **(Figures S3C & S4C)**. Although the MAE between the posterior means and the assigned interaction coefficients decreased slightly with 6 observations, it slightly increased with 8 and 10 observations, possibly due to minor discrepancies in numerical integrations at these intermediate time points. In contrast, using 25 observations consistently produced the lowest MAE **(Figure 3D)**. While increasing the number of observations reduced the uncertainty of parameter estimates, gains in accuracy plateaued beyond a certain point, with additional observations offering little improvement.

**Figure 3:**
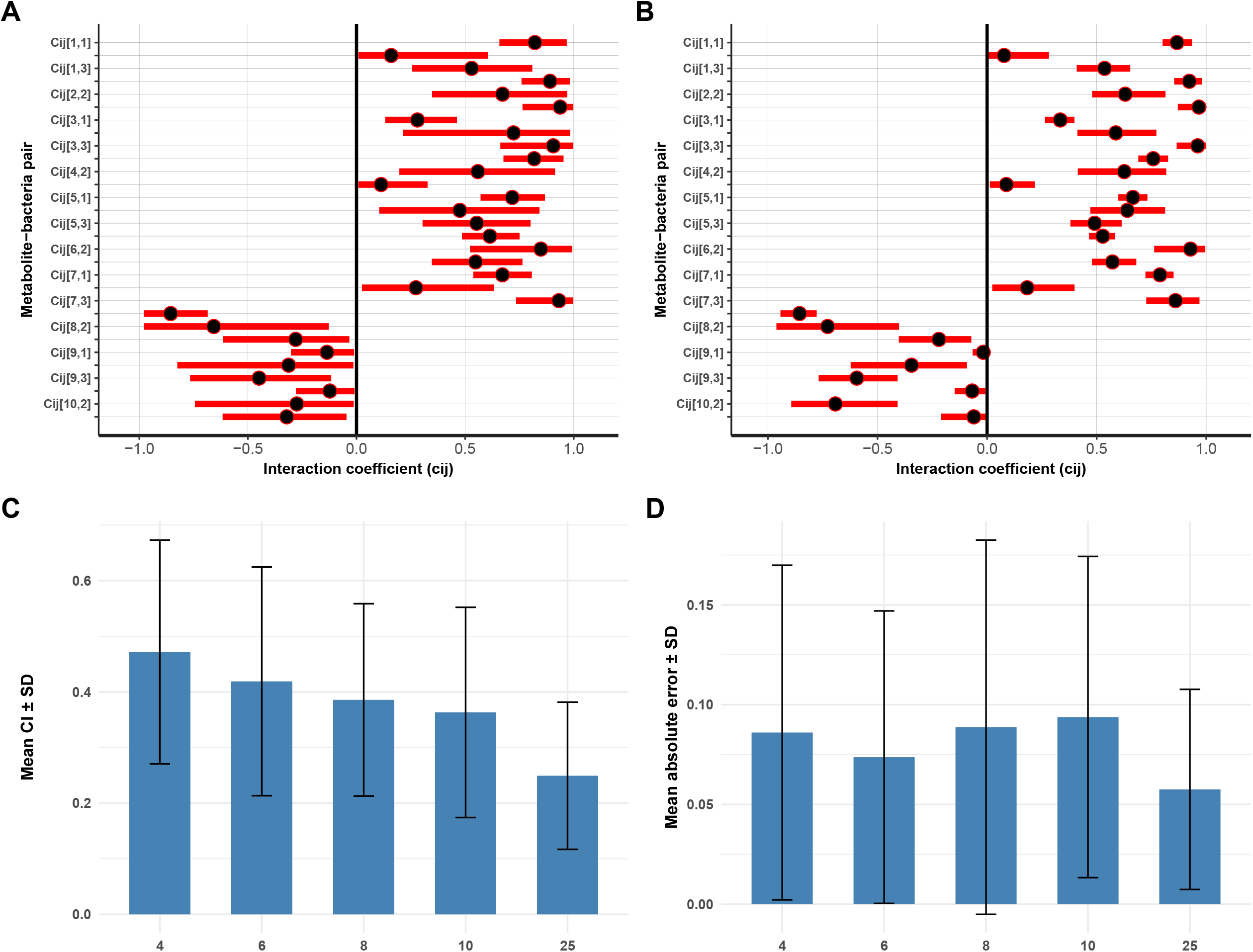
Estimation uncertainty increases with observation error but decreases with more observations. A) Posterior distributions of all interaction coefficients (*c*_*i*,*j*_) in the small bacterial community using four observations with observation error added. Posterior means are shown as black dots and 95% credible intervals as red bars. B) Posterior distributions of all interaction coefficients in the same community using 25 observations with observation error added. Posterior means are shown as black dots and 95% credible intervals as red bars. C) Mean and standard deviation (SD) of 95% credible intervals (CI) across posterior distributions obtained from parameter estimation with varying numbers of observations in the small bacterial community with added observation error. D) Mean and SD of the absolute error between assigned interaction coefficients and posterior means obtained from parameter estimation with varying numbers of observations in the small bacterial community with added observation error.

### Accuracy declines with community size

We only considered a small bacterial system with three bacteria and ten metabolites, based on a synthetic community studied in wet lab experiments [21]. However, many synthetic communities contain more species and metabolites [23], and naturally occurring communities can be even larger and more complex [24]. To assess the feasibility and accuracy of our approach in larger communities, we applied Bayesian inference to ten medium-sized communities consisting of 5 bacteria and 14 metabolites, and to five large communities consisting of 10 bacteria and 20 metabolites. Model diagnostics for both community sizes indicated no divergent transitions, with Rhat values close to one and large `n_eff` **(Figures S5A & B; S6A & B)**. Comparisons of posterior distributions **(Figures S5C & S6C)**, posterior predictive checks **(Figures S5D & S6D)**, and comparison of the assigned interaction coefficients with corresponding posterior means **(Figures S5E & 6E)** showed consistent patterns across communities. Specifically, as community size increased, posterior means for consumed metabolites shifted towards 0, those for produced metabolites shifted towards -0.5, and the posteriors became flatter, indicating reduced predictive power. Furthermore, the credible intervals of the posterior distributions **(Figure 4A)** and the MAE between the assigned interaction coefficients and corresponding posterior means **(Figure 4B)**, both increased with community size, indicating reduced accuracy and reliability of Bayesian inference in larger communities.

**Figure 4:**
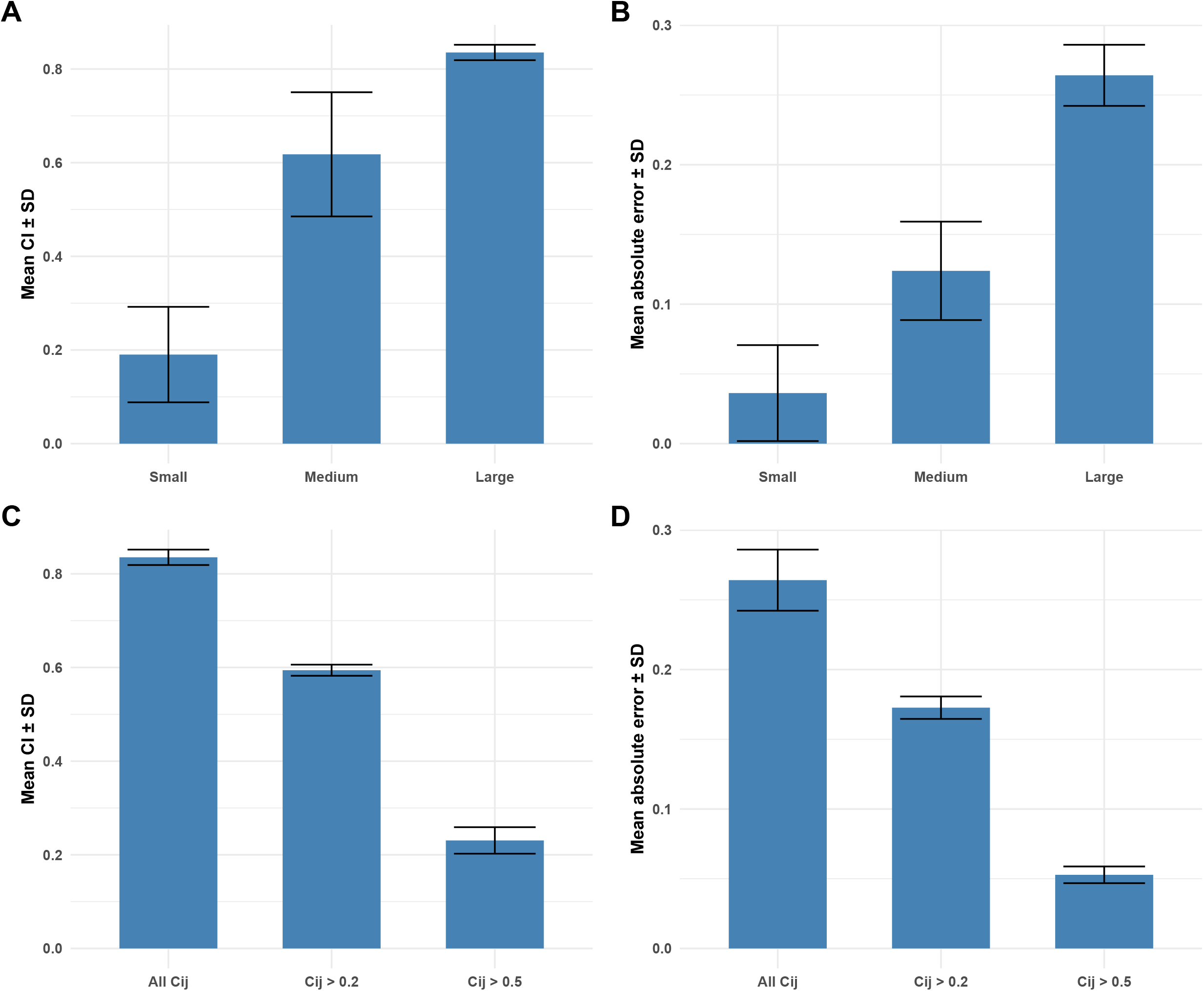
Estimation uncertainty and accuracy vary with community size and network sparsity. A) Mean and standard deviation (SD) of the 95% credible intervals (CI) across posterior distributions obtained from parameter estimation using four observations for three community sizes (small: 3 bacteria and 10 metabolites; medium: 5 bacteria and 14 metabolites; large: 10 bacteria and 20 metabolites). B) Mean and standard deviation (SD) of the absolute error between assigned interaction coefficients and posterior means for the same three community sizes. C) Mean and SD of the 95% CI across posterior distributions for the large community (10 bacteria and 20 metabolites) under three sparsity conditions: all interaction coefficients included (All *c*_*i*,*j*_), coefficients below 0.2 removed (*c*_*i*,*j*_ > 0.2), and coefficients below 0.5 removed (*c*_*i*,*j*_ > 0.5). D) Mean and SD of the absolute error between assigned interaction coefficients and posterior means for the large community under the same sparsity conditions as in panel C.

### Imposing sparsity of the interaction matrix improves estimation

In the analyses above, an interaction coefficient was generated for every bacterium-metabolite pair. However, in both synthetic and natural communities, not every metabolite is produced or consumed by every bacterium; i.e., the bipartite network of bacterium-metabolite interactions is not fully connected. In our simulations, we observed that as community size increased, posterior distributions for interactions between bacteria and consumed metabolites concentrated near zero. This suggests that certain metabolites do not contribute to the growth of specific bacteria and could be omitted in simulated communities.

To investigate whether reducing the number of non-zero interaction coefficients improved model accuracy, we discarded interaction coefficients with absolute values below 0.2 and 0.5 in the large community, eliminating 38 (19%) and 114 (57%) interaction coefficients, respectively. Using the same inference approach, model diagnostics for both sparser communities showed no divergent transitions, with `Rhat` values close to one and large `n_eff` **(Figures S7A & B; S8A & B)**. Comparisons of posterior distributions **(Figures S7C & S8C)**, posterior predictive checks **(Figures S7D & S8D)**, and comparison of assigned interaction coefficients with corresponding posterior means **(Figures S7E & S8E)** showed that the posteriors became more concentrated, with clearer modes moving away from zero as the number of non-zero interaction coefficients decreased. Credible intervals narrowed substantially in the sparser communities **(Figure 4C)**, and the MAE between the assigned interaction coefficients and corresponding posterior means also declined **(Figure 4D)**. These results indicate that Bayesian inference can identify interaction structures in large communities, provided that the connectance of the microbial network (i.e., the number of non-zero interaction coefficients) is sufficiently low.

## Discussion

A lack of theoretical foundation is hampering our understanding of the structure and function of microbial communities and consequently limits the development of novel microbiota-targeted interventions to improve the health of macro-ecosystems. Central to this problem is to infer and quantify microbial interactions and to develop predictive tools for community dynamics. Here, we developed a Bayesian inference framework that is able to accurately and reliably quantify interactions between bacteria and metabolites in simulated microbial communities. In small communities, we have shown that with noisy data due to observation errors, increasing the number of observations improves the accuracy of parameter estimation. In larger communities, our approach remains viable provided that the interaction network is relatively sparse. Overall, this framework provides a way to characterise microbial communities and lays the foundation for designing microbiota-targeted interventions to support macro-ecosystem health.

### Generative models with diverse growth forms

A wide variety of quantitative methods have been used to capture microbial interactions, including Lotka-Volterra type models [25], correlation-based approaches [26], consumer-resource models [18] and agent-based models [27]. Modelling microbial interactions with these quantitative methods facilitates our understanding of the structure, dynamics and functional properties of microbial networks [28]. Our generative model quantifies interactions using a set of coupled ODEs that describe the dynamics of consuming and excreting metabolites in a bacterial community. The growth of each bacterial species is described by the sum of all consumed metabolites multiplied by their corresponding interaction coefficients, thereby representing substitutable and additive bacterial growth dynamics commonly employed in consumer-resource models of microbial communities [18, 29]. This type of growth dynamics has been shown to predict microbial composition at the family level with high accuracy in an experimental setup [30]. Nonetheless, for specific systems or metabolic environments, more intricate forms of the per-capita growth function *g*_*j*_(***M***) may be required. For example, even under the assumption of substitutable metabolites, per-capita growth *g*_*j*_(***M***) saturates when metabolite concentrations are high. Because uptake and conversion rates are physiologically constrained by transporter and enzyme capacities, growth dynamics are better described by Monod kinetics: 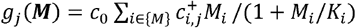, with *K*_*i*_ the half-saturation constant [31, 32]. The per-capita growth can also be limited by the scarcest metabolite following Liebig’s minimum principle, 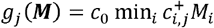, or be co-limited by multiple metabolites [33], 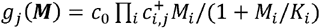. Besides substitutable, some metabolites such as toxins can inhibit growth, 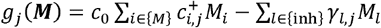, with {inh} the set of inhibitory metabolites and *γ*_*l*,*j*_ the strength of inhibition of metabolite *l* on bacterium *j*. More complex growth forms can also be considered, e.g., bacterium *j* does not uptake a metabolite for growth until the concentration has reached a threshold [34]. All these variants of *g*_*j*_(***M***) can be implemented in Eq. (1), depending on the research question or specific system of interest. To account for more complex metabolite dynamics such as overflow metabolism, cross feeding and other non-monotonic behaviours, Hill-type functions can be incorporated into Eq. (2) [35]. In our simulations, the carrying capacity, metabolite conversion coefficients and death rates were held constant. These parameters can also be adjusted to better reflect the community under study or estimated using the same Bayesian inference framework by assigning appropriate priors, though increasing model complexity inevitably raises computational demands.

### Bayesian inference with informative priors

Prior knowledge of the system under study can be incorporated to add biological realism. Adjusting specific interaction coefficients is particularly valuable, as constraining more entries of the interaction matrix to zero improves estimation for the remaining coefficients **(Figure 4C and D)**. For example, in a simple two-species community, *Bifidobacterium adolescentis* produces ethanol and acetate, while *Faecalibacterium prausnitzii* consumes acetate and produces butyrate [36]. In this case, the interaction coefficient between *F. prausnitzii* and ethanol, and between *B. adolescentis* and butyrate, can be set to zero, while the signs of the interaction between both bacteria and acetate can be fixed as well. Genome-scale metabolic models (GEMs) are useful tools to obtain such prior knowledge as these models are constructed from the genomic sequences and consist of experimentally validated or predicted biochemical reactions [37]. While GEMs for well-studied bacteria and communities (e.g. the human gut) are generally accurate and reliable, those for rare or poorly studied bacteria often contain erroneous or incomplete biochemical reactions [38].

Beyond adjusting specific parameters, such as metabolite conversion rates, bacterial growth forms or interaction coefficients, Bayesian statistical methods offer an inherent way to integrate biological realism. Model assumptions are made explicit through the choice of likelihoods and priors. Estimation accuracy can be improved by optimising prior distributions or embedding prior selection in a reinforcement learning framework, where updated posteriors serve as priors as new data become available. In our study, posterior distributions of interaction coefficients were derived using broad uniform priors; narrowing these intervals or using alternative priors may better reflect biological reality and improve inference [39]. For the likelihood function, we used a lognormal distribution, constraining outcomes to non-negative, skewed distributions with multiplicative variability. This fits well with microbial systems: since metabolite concentrations and bacterial counts cannot be negative, most microbial communities follow highly skewed abundance distributions [40], and small differences in enzyme expression and catalysis can compound over time, affecting abundance distributions of bacterial taxa [41]. Environmental perturbations can shift species abundance patterns and alter interaction networks [42] as observed when comparing bacteria-metabolite networks from healthy versus type-2 diabetic gut microbiota [43], potentially requiring alternative likelihood choices.

Finally, Bayesian inference regularizes parameter estimation due to the use of priors which reduces overfitting even with small sample sizes or relatively few measurements [17]. While additional observations did improve accuracy of parameter estimation in our approach, the effect was modest **(Figure 3)**. Thus, Bayesian methods are particularly suitable for studying dynamical microbial systems, where sampling is often constrained by costs and logistics [44, 45]. Moreover, Bayesian methods explicitly quantify uncertainty, accommodating both stochasticity and experimental noise. Posterior distributions can be propagated to generate predictive distributions of microbial dynamics or metabolite trajectories, offering a direct means to assess responses to perturbations and to design microbiota-targeted interventions.

### Concluding remarks

The proposed Bayesian inference framework provides an accurate and reliable means of estimating bacteria-metabolite interactions in simulated microbial networks. The generative model offers a flexible backbone for representing diverse microbial communities, while its numerical implementation and Bayesian inference enable the quantification of network structures from empirical bacterial and metabolic data. When applied to real-world microbial communities, this approach can serve as a means to simulate the responses of microbiota-targeted interventions *in-silico* by altering system properties such as network connectance, interaction strengths and species composition. In doing so, it can help identify possible intervention targets, including probiotics, prebiotics and dietary alterations. Our combined approach may guide the development of microbiota-targeted interventions to improve the health of macro-ecosystems, including the human gut, by establishing a theoretical foundation for future studies.

## Data availability

All code used in this study is publicly available at: https://github.com/JackJansma/Bayesian-inference-of-bacteria-metabolite-interactions. This includes R scripts for: (i) simulating bacteria–metabolite dynamics using ODE models, (ii) performing Bayesian parameter inference, and (iii) processing and visualizing the results. The simulations and analyses can be fully reproduced using the provided scripts, with the specified random seed ensuring reproducibility. The repository also includes documentation and information about required R packages and software dependencies. The generated data regarding all simulated communities and all parameter estimations using the Bayesian inference framework is publicly available at: https://doi.org/10.5281/zenodo.17078524

## Author contributions

J.J., P.L and C.H. conceived and designed the study (Conceptualization, Methodology). J.J. performed the simulations and analysed the data (Software, Formal analysis). J.J. drafted the manuscript (Writing – original draft). All authors contributed to reviewing and editing the manuscript (Writing – review & editing) and approved the final version.

