## Supplementary file for "Bayesian inference captures metabolite-bacteria interactions in a microbial community"

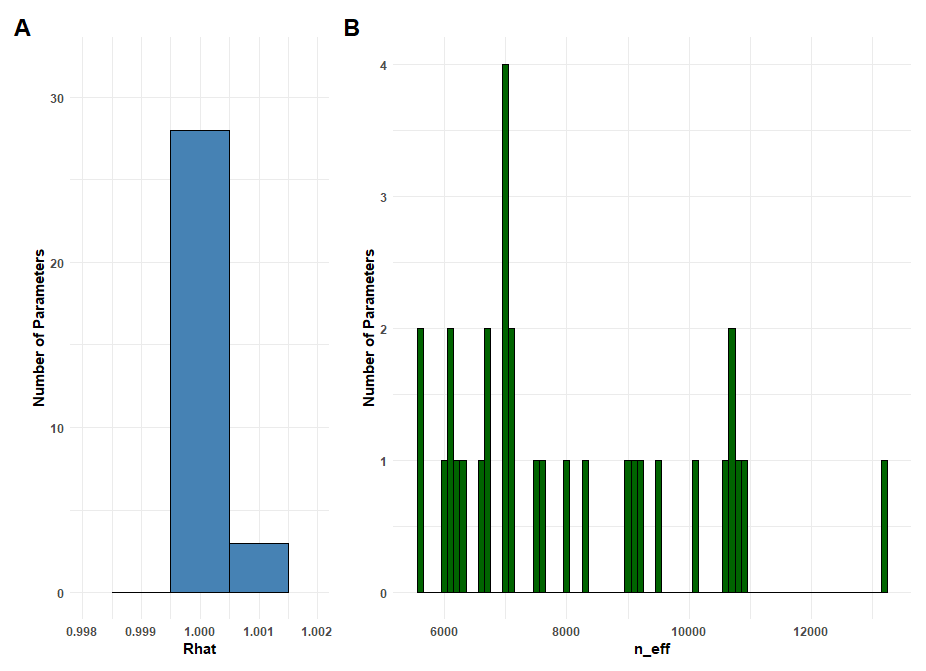

**Supplementary figure 1: Model diagnostics of parameter estimation in a small bacterial community.**

A) Rhat values for all estimated interaction coefficients. B) Effective sample sizes (n_eff) for all estimated interaction coefficients. Rhat and n_eff values are obtained from 1 representative example out of the total of 250 model fits. Model fitting was conducted by running four chains for 2000 iterations, with the first 1000 iterations discarded as warm-up.

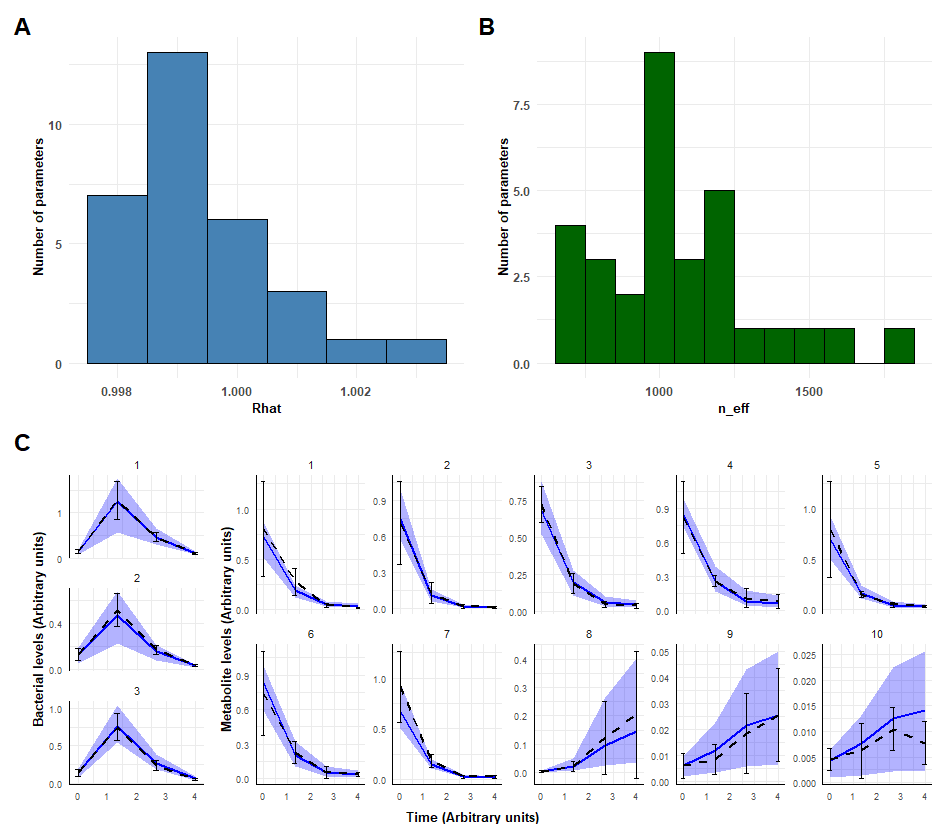

**Supplementary figure 2: Model diagnostics and posterior predictive check of parameter estimation in a small bacterial community with noise addition.**

A) Rhat values for all estimated interaction coefficients. B) Effective sample sizes (n_eff) for all estimated interaction coefficients. Model fitting was conducted by running two chains for 1000 iterations, with the first 500 iterations discarded as warm-up. C) Posterior predictive check. The mean and standard deviation of four generated observations with added noise, obtained by numerically solving the set of ODEs using the assigned initial conditions and interaction coefficients, are shown as black dotted lines and error bars. The mean and 95% credible intervals of generated observations, using draws from the estimated posterior distributions, are shown as a blue line and a purple belt.

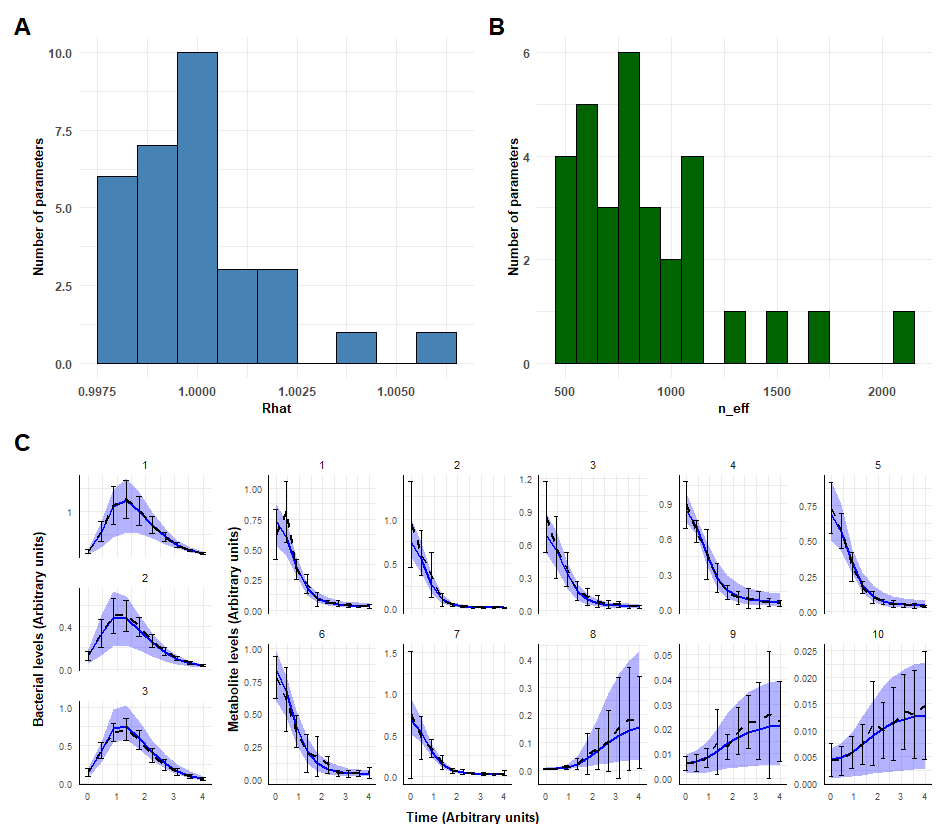

**Supplementary figure 3: Model diagnostics and posterior predictive check of parameter estimation in a small bacterial community with noise addition and ten observations.**

A) Rhat values for all estimated interaction coefficients. B) Effective sample sizes (n_eff) for all estimated interaction coefficients. Model fitting was conducted by running two chains for 1000 iterations, with the first 500 iterations discarded as warm-up. C) Posterior predictive check. The mean and standard deviation of ten generated observations with added noise, obtained by numerically solving the set of ODEs using the assigned initial conditions and interaction coefficients, are shown as black dotted lines and error bars. The mean and 95% credible intervals of generated observations, using draws from the estimated posterior distributions, are shown as a blue line and a purple belt.

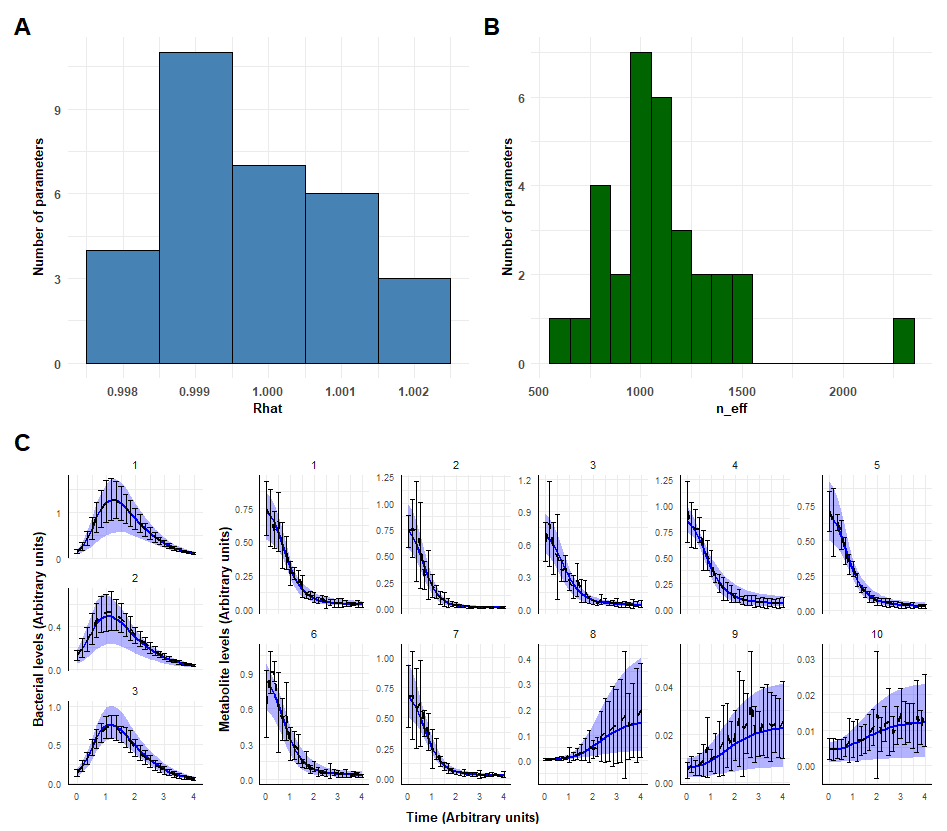

**Supplementary figure 4: Model diagnostics and posterior predictive check of parameter estimation in a small bacterial community with noise addition and 25 observations.**

A) Rhat values for all estimated interaction coefficients. B) Effective sample sizes (n_eff) for all estimated interaction coefficients. Model fitting was conducted by running two chains for 1000 iterations, with the first 500 iterations discarded as warm-up. C) Posterior predictive check. The mean and standard deviation of 25 generated observations with added noise, obtained by numerically solving the set of ODEs using the assigned initial conditions and interaction coefficients, are shown as black dotted lines and error bars. The mean and 95% credible intervals of generated observations, using draws from the estimated posterior distributions, are shown as a blue line and a purple belt.

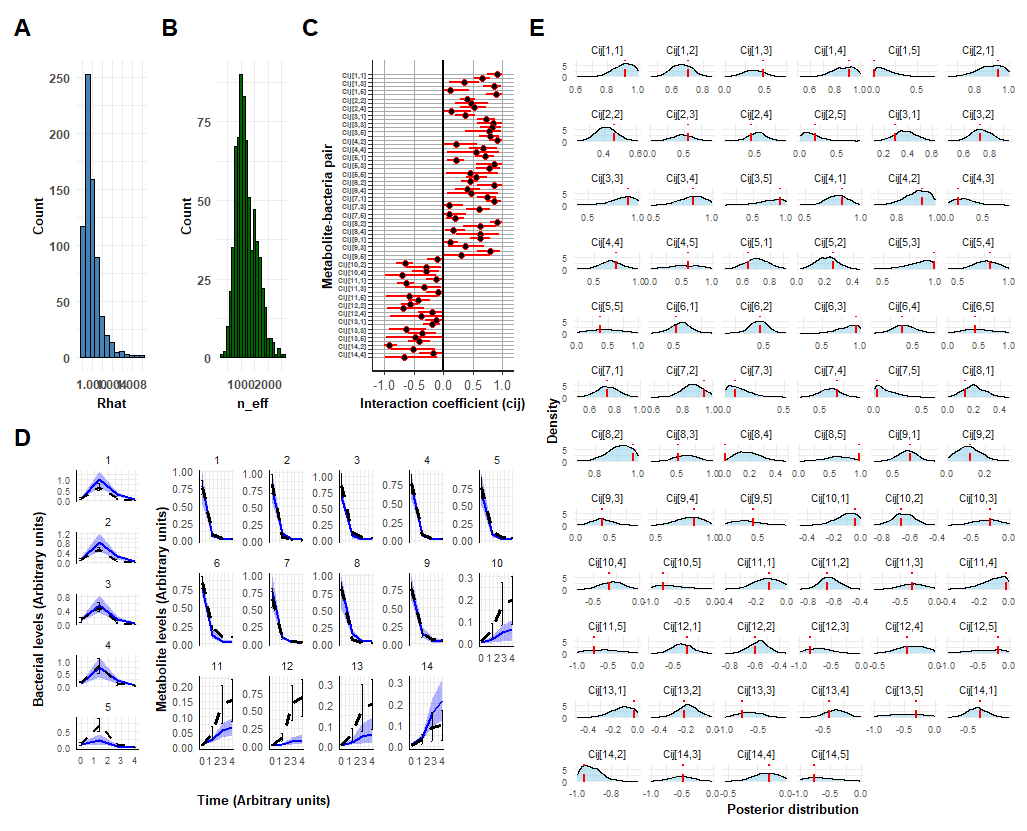

**Supplementary figure 5: Overview of the parameter estimation in a medium sized bacterial community consisting of 5 bacteria and 14 metabolites.**

A) Rhat values for all estimated interaction coefficients. B) Effective sample sizes (n_eff) for all estimated interaction coefficients. Model fitting was conducted by running two chains for 1000 iterations, with the first 500 iterations discarded as warm-up. C) Posterior distributions of all interaction coefficients, $c_{i,j}$. Posterior means are depicted as black dots, with 95% credible intervals indicated by red bars. D) Posterior predictive check. The mean and standard deviation of four generated observations, obtained by numerically solving the set of ODEs using the assigned initial conditions and interaction coefficients, are shown as black dotted lines and error bars. The mean and 95% credible intervals of generated observations, using draws from the estimated posterior distributions, are shown as a blue line and a purple belt. E) Comparisons of assigned interaction coefficients (red dotted lines) with the corresponding posterior distributions. All plots show the results of 1 representative example out of a total of 10 model fits.

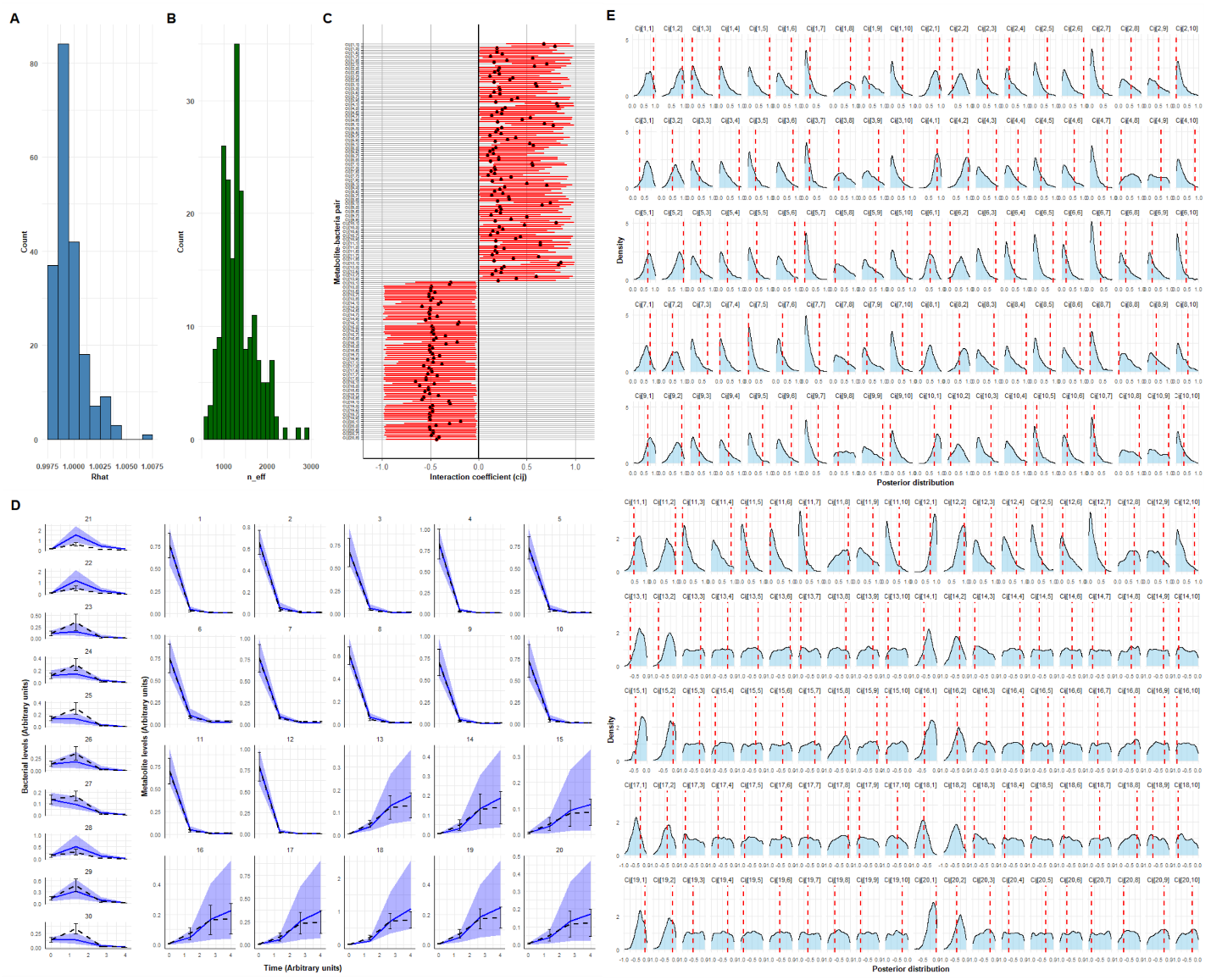

**Supplementary figure 6: Overview of the parameter estimation in a large bacterial community consisting of 10 bacteria and 20 metabolites.**

A) Rhat values for all estimated interaction coefficients. B) Effective sample sizes (n_eff) for all estimated interaction coefficients. Model fitting was conducted by running two chains for 1000 iterations, with the first 500 iterations discarded as warm-up. C) Posterior distributions of all interaction coefficients, $c_{i,j}$. Posterior means are depicted as black dots, with 95% credible intervals indicated by red bars. D) Posterior predictive check. The mean and standard deviation of four generated observations, obtained by numerically solving the set of ODEs using the assigned initial conditions and interaction coefficients, are shown as black dotted lines and error bars. The mean and 95% credible intervals of generated observations, using draws from the estimated posterior distributions, are shown as a blue line and a purple belt. E) Comparisons of assigned interaction coefficients (red dotted lines) with the corresponding posterior distributions. All plots show the results of 1 representative example out of a total of 10 model fits.

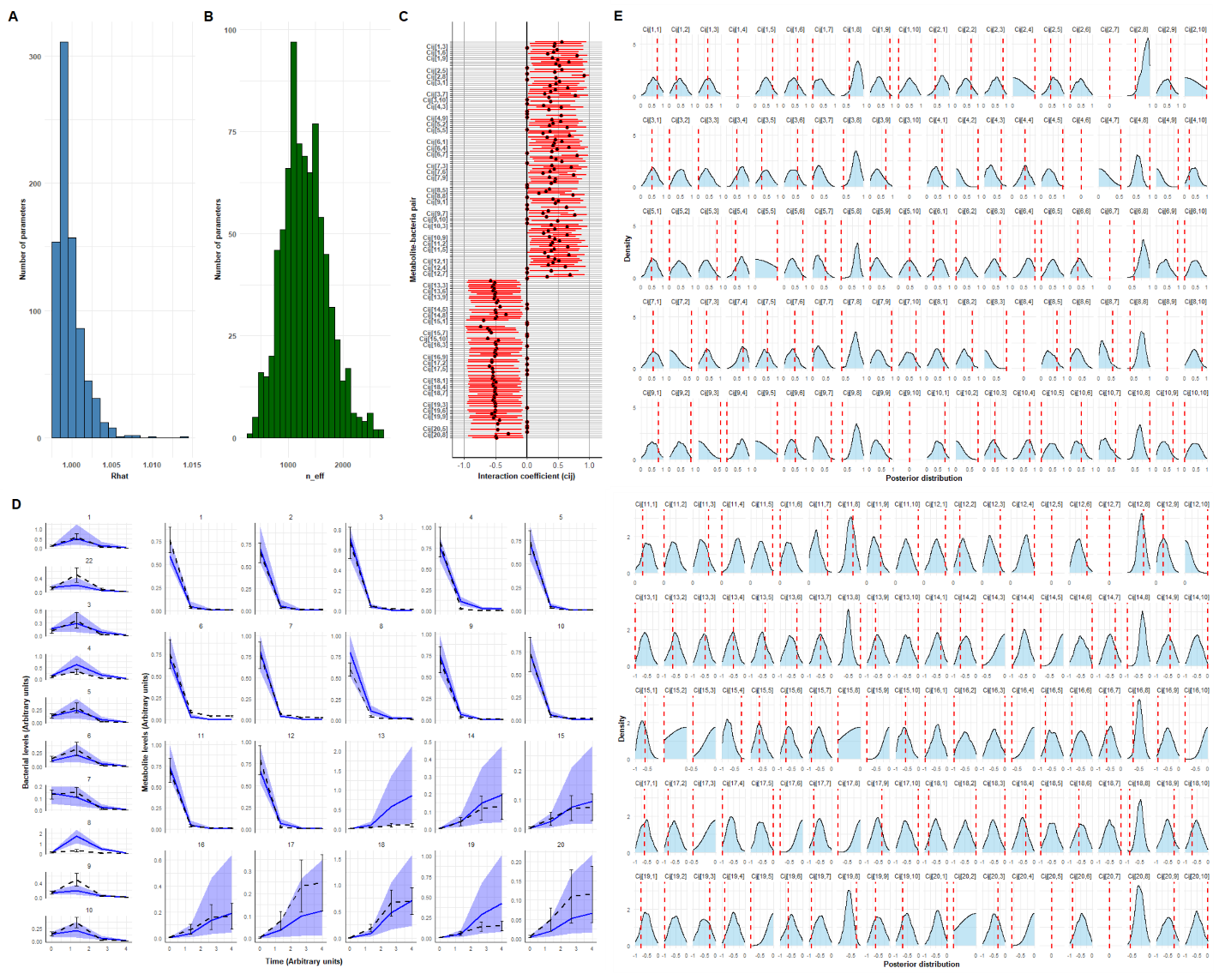

**Supplementary figure 7: Overview of the parameter estimation in a large bacterial community without interaction coefficients below 0.2.**

A) Rhat values for all estimated interaction coefficients. B) Effective sample sizes (n_eff) for all estimated interaction coefficients. Model fitting was conducted by running two chains for 1000 iterations, with the first 500 iterations discarded as warm-up. C) Posterior distributions of all interaction coefficients, $c_{i,j}$. Posterior means are depicted as black dots, with 95% credible intervals indicated by red bars. D) Posterior predictive check. The mean and standard deviation of four generated observations, obtained by numerically solving the set of ODEs using the assigned initial conditions and interaction coefficients, are shown as black dotted lines and error bars. The mean and 95% credible intervals of generated observations, using draws from the estimated posterior distributions, are shown as a blue line and a purple belt. E) Comparisons of assigned interaction coefficients (red dotted lines) with the corresponding posterior distributions. All plots show the results of 1 representative example out of a total of 5 model fits.

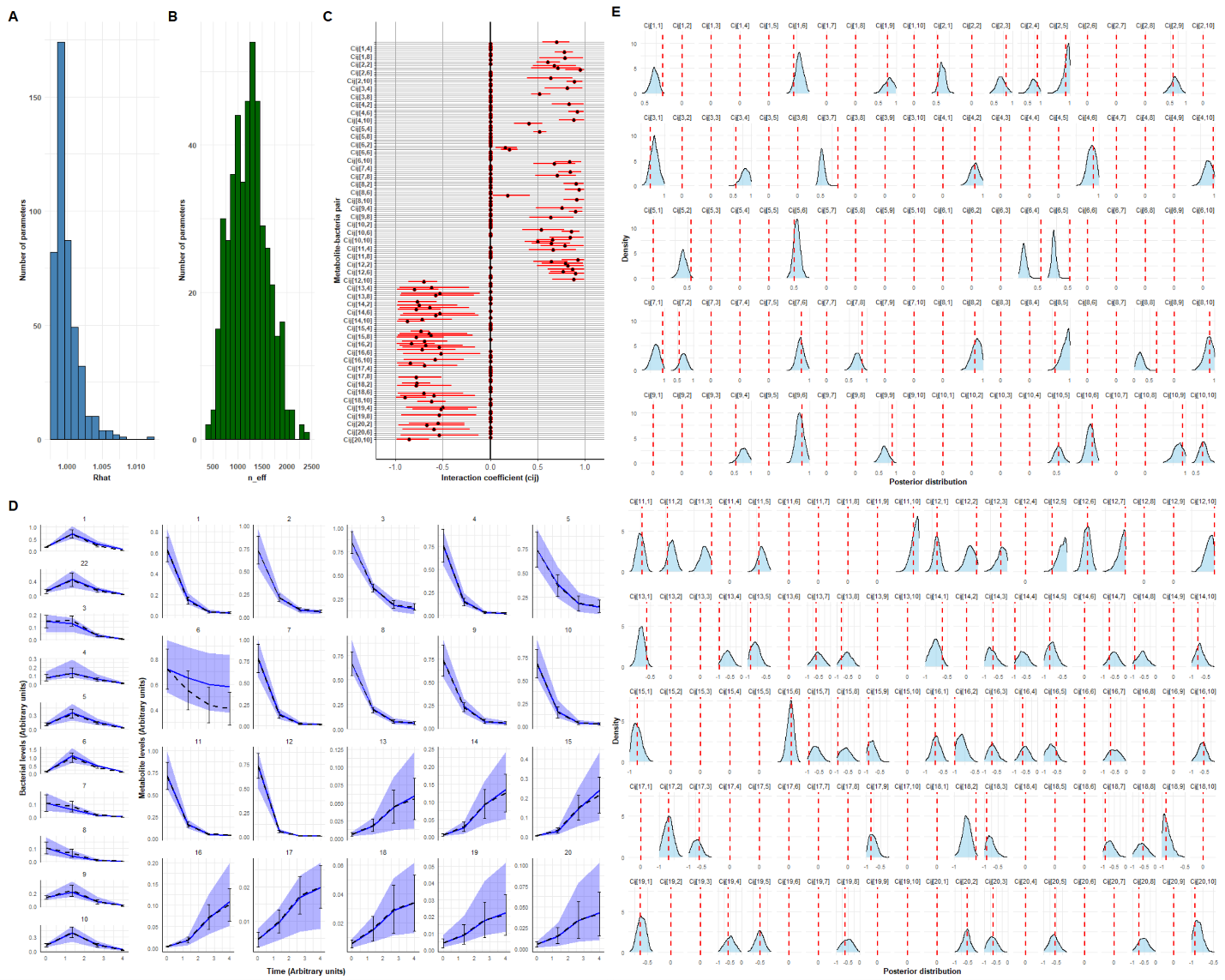

**Supplementary figure 8: Overview of the parameter estimation in a large bacterial community without interaction coefficients below 0.5.**

A) Rhat values for all estimated interaction coefficients. B) Effective sample sizes (n_eff) for all estimated interaction coefficients. Model fitting was conducted by running two chains for 1000 iterations, with the first 500 iterations discarded as warm-up. C Posterior distributions of all interaction coefficients, $c_{i,j}$. Posterior means are depicted as black dots, with 95% credible intervals indicated by red bars. D) Posterior predictive check. The mean and standard deviation of four generated observations, obtained by numerically solving the set of ODEs using the assigned initial conditions and interaction coefficients, are shown as black dotted lines and error bars. The mean and 95% credible intervals of generated observations, using draws from the estimated posterior distributions, are shown as a blue line and a purple belt. E) Comparisons of assigned interaction coefficients (red dotted lines) with the corresponding posterior distributions. All plots show the results of 1 representative example out of a total of 5 model fits.
